# LARP1 facilitates translational recovery after amino acid refeeding by preserving long poly(A)-tailed TOP mRNAs

**DOI:** 10.1101/716217

**Authors:** Koichi Ogami, Yuka Oishi, Takuto Nogimori, Kentaro Sakamoto, Shin-ichi Hoshino

## Abstract

Occasionally, cells must adapt to an inimical growth conditions like amino acid starvation (AAS) by downregulating protein synthesis. A class of transcripts containing 5’terminal oligopyrimidine (5’TOP) motif encodes translation-related proteins such as ribosomal proteins (RPs) and elongation factors, and therefore, their translation is severely repressed during AAS to conserve energy^1^. The RNA-binding protein LARP1 transduces amino acid signaling to TOP gene expression by controlling translation and stability of TOP mRNAs^2-6^. When released from AAS, translation machineries in turn have to be restored, however, the underlying mechanism of such re-adaptation is largely unknown. Here we show that LARP1 preserves TOP mRNAs in a long polyadenylated state during long-term AAS. We found that TOP mRNAs become highly polyadenylated when cells are in AAS or treated with the mTOR (mechanistic target of rapamycin) inhibitor Torin1. Importantly, depletion of LARP1 completely abrogated the polyadenylation of TOP mRNAs. Comprehensive analysis of poly(A) tail length using the Nanopore direct RNA sequencing revealed that TOP mRNAs are selectively polyadenylated under mTOR inhibition. Since a long poly(A) tail confers increased stability and polysome formation of TOP mRNAs, we predict that LARP1-dependent preservation of TOP mRNAs enables rapid translational resumption after the release from AAS.

## Results and Discussion

We first investigated the effect of long-term AAS (∼12 h) on three RP mRNAs (RPS6, RPL11 and RPL26 mRNA). Northern blotting revealed that mRNA species with slower migration rate started to accumulate constantly during AAS, while the fast migrating species that are major in normal culture condition (before starvation; 0 h) gradually declined (Fig. 1a). Poly(A) tail removal by oligo(dT)/RNase H digestion rendered a single band at the same position, indicating that these mRNAs are highly polyadenylated during the prolonged starvation (Fig. 1a). Poly(A) tail length of these mRNAs are dynamically regulated in response to amino acid availability since the elongated tails were shortened again when starved cells were re-fed (Fig. 1b). Accumulation of long-tailed RP mRNAs was also observed when cells were treated with Torin1 (Fig. 1c), indicating that poly(A) elongation occurs downstream of the amino acid-mTOR signaling pathway. In contrast to RP mRNAs, tail length of a nonTOP transcript, GAPDH mRNA, is shortened during AAS and Torin1 treatment (Fig. 1a-c). To analyze poly(A) tail length independently from transcription, we employed the 4-thiouridine (4SU) labeling method (Fig. 1d). Poly(A) tail of the labeled RP mRNAs remained long in starved or Torin1-treated cells (-AA, Torin1), which is in contrast to the progressive poly(A) shortening in the normal culture condition (Fed) (Fig. 1e). Consistent results were obtained using transcription inhibition using actinomycin D (ActD): poly(A) tail shortening of RP mRNAs is impaired by Torin1 treatment (Fig. 1f). LARP1 directly interacts with 5’TOP motifs and the adjacent cap structure of RP mRNAs, especially during AAS and mTOR inhibition^4,7-9^. LARP1 also interacts with 3’poly(A) tail directly^10^ as well as through the poly(A)-binding protein PABPC1^5,10,11^, although the role of 3’ terminal association remains unknown. Strikingly, AAS- or Torin1-induced tail elongation was severely abolished by LARP1 knockdown (KD) using two different siRNAs (Fig.1g-i).

**Figure 1.**
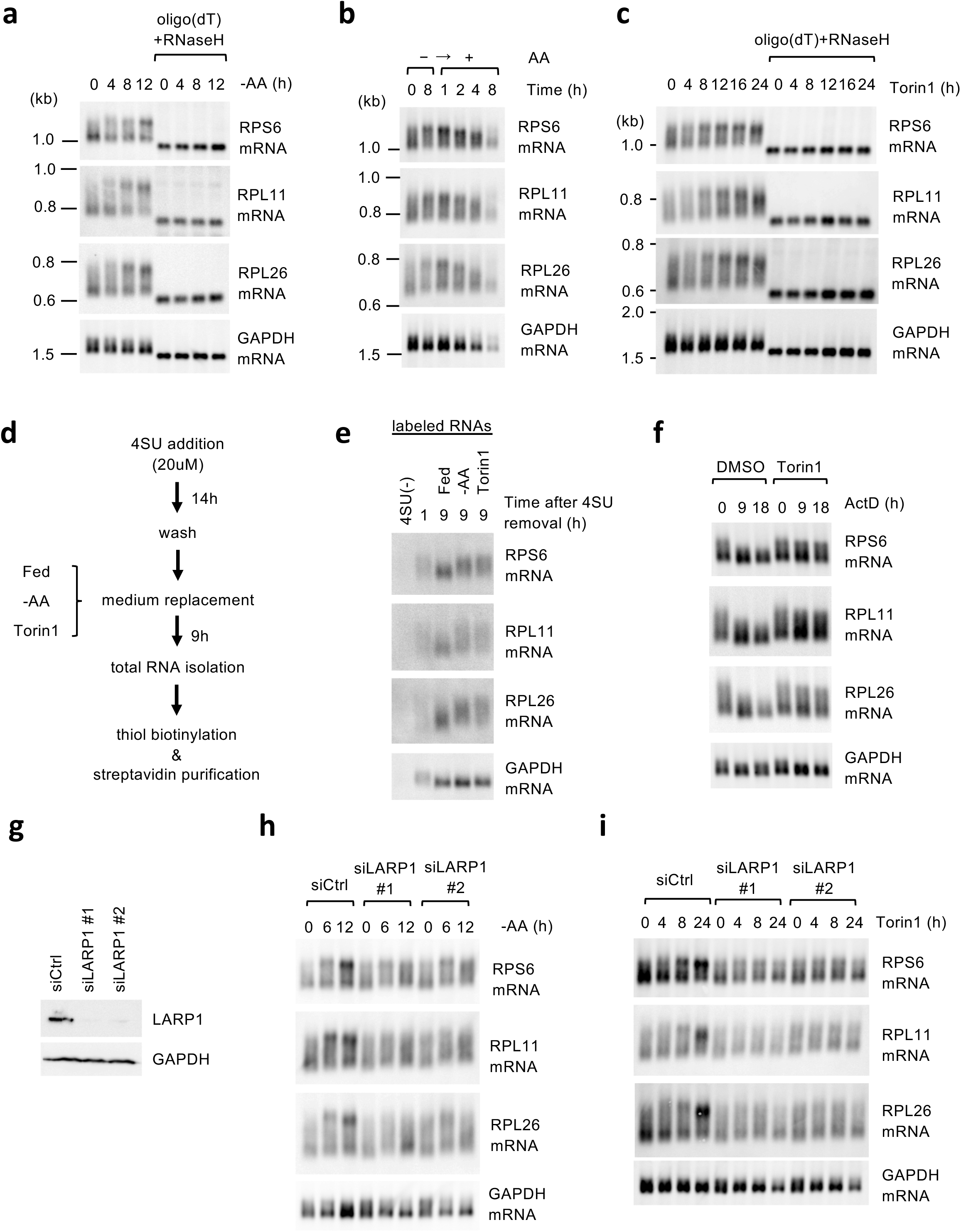
TOP mRNA with a long poly(A) tail accumulates after amino acid starvation in a LARP1-dependent manner. **a**, Northern blotting of total RNA extracted from HEK293T cells cultured in the medium without amino acid (-AA) for the indicated times. Poly(A) removal was performed by oligo(dT)/RNaseH treatment of the total RNA. **b**, Amino acid-starved HEK293T cells were re-fed with complete DMEM. Total RNA was isolated at the indicated time points and analyzed by Northern blotting. **c**, Total RNA from HEK293T cells treated with 250 nM Torin1 in complete DMEM for the indicated times was analyzed by Northern blotting. Poly(A) removal was performed by oligo(dT)/RNaseH treatment of the total RNA. **d**, Scheme of the 4SU labeling experiments. After labeling with 4SU for 14 hrs, HEK293T cells were washed and cultured for additional 9 hrs in 4SU-free medium under Fed, -AA or 250 nM Torin1 conditions. The labeled RNAs were thiol-biotinylated and purified using streptavidin beads. **e**, The labeled RNAs purified as in **d** were analyzed by Northern blotting. **f**, HEK293T cells were treated with 10 μ g/mL actinomycin D together with 250 nM Torin1, and total RNAs isolated at the indicated time points were analyzed by Northern blotting. **g**, Western blotting showing the knockdown efficiency after 48 hrs of siRNA transfection in HEK293T cells. **h, i**, After siRNA transfection, HEK293T cells were cultured in DMEM without amino acids (-AA) (**h**), complete DMEM containing 250 nM Torin1 (**i**). Total RNAs were isolated at the indicated time points and analyzed by Northern blotting.

To gain a whole picture of the poly(A) tail length changes under mTOR inhibition as well as LARP1 KD, we set out to conduct a high-throughput poly(A) tail analysis of the intact mRNA using the nanopore direct RNA sequencing (dRNA-seq)^12^. dRNA-seq obtains long sequence reads from native RNAs and therefore provides an advantage over next generation sequencing (NGS)-based approaches: dRNA-seq eliminates PCR amplification biases that can lead to over- and under-representation of certain sequences including poly(A) tails. To enrich mRNAs, we employed a 5’cap-purification strategy using an eIF4E K119A with high-affinity for m^7^G-cap (Supplementary Fig. 1a)^13^. We first removed highly abundant 5’ capped small non-coding RNAs (ncRNAs) from the extracted total RNA by repeating LiCl precipitation twice to avoid competition for eIF4E_K119A_-binding (Supplementary Fig. 2b). 5’capped mRNAs were then collected using GST-eIF4E_K119A_. Sequencing libraries were prepared following the Oxford nanopore’s instruction for dRNA-seq. Notably, as 5’cap-purified RNAs were ligated with the double-stranded adapter with oligo(dT)_10_ overhang that anneals to the 3’-most end of a poly(A) tail, the final RNA libraries were all originated from the intact non-truncated transcripts (Fig. 2a). We obtained 1.1∼1.3 million reads (total bases of ∼1.6 billion) with good average quality scores of Q ∼9 (Supplementary Fig. 1c). Poly(A) tail length of the each read was estimated using nanopolish polya program^14^. Without Torin1 treatment, density plots of TOP mRNA show that tail length distribution is largely right-skewed with a peak around 50 nt (siCtrl and siLARP1), while siCtrl-Torin1 exhibited bimodal distribution of TOP mRNA with a secondary peak around 150 nt and these changes were cancelled by LARP1 KD (siLARP1-Torin1) (Fig. 2a). Consistent results were obtained in the tail length distribution of individual TOP transcripts (Fig. 2b). On the contrary, such tail length changes were not apparent in nonTOP mRNAs (Fig. 2a, Supplementary Fig. 2).

**Figure 2.**
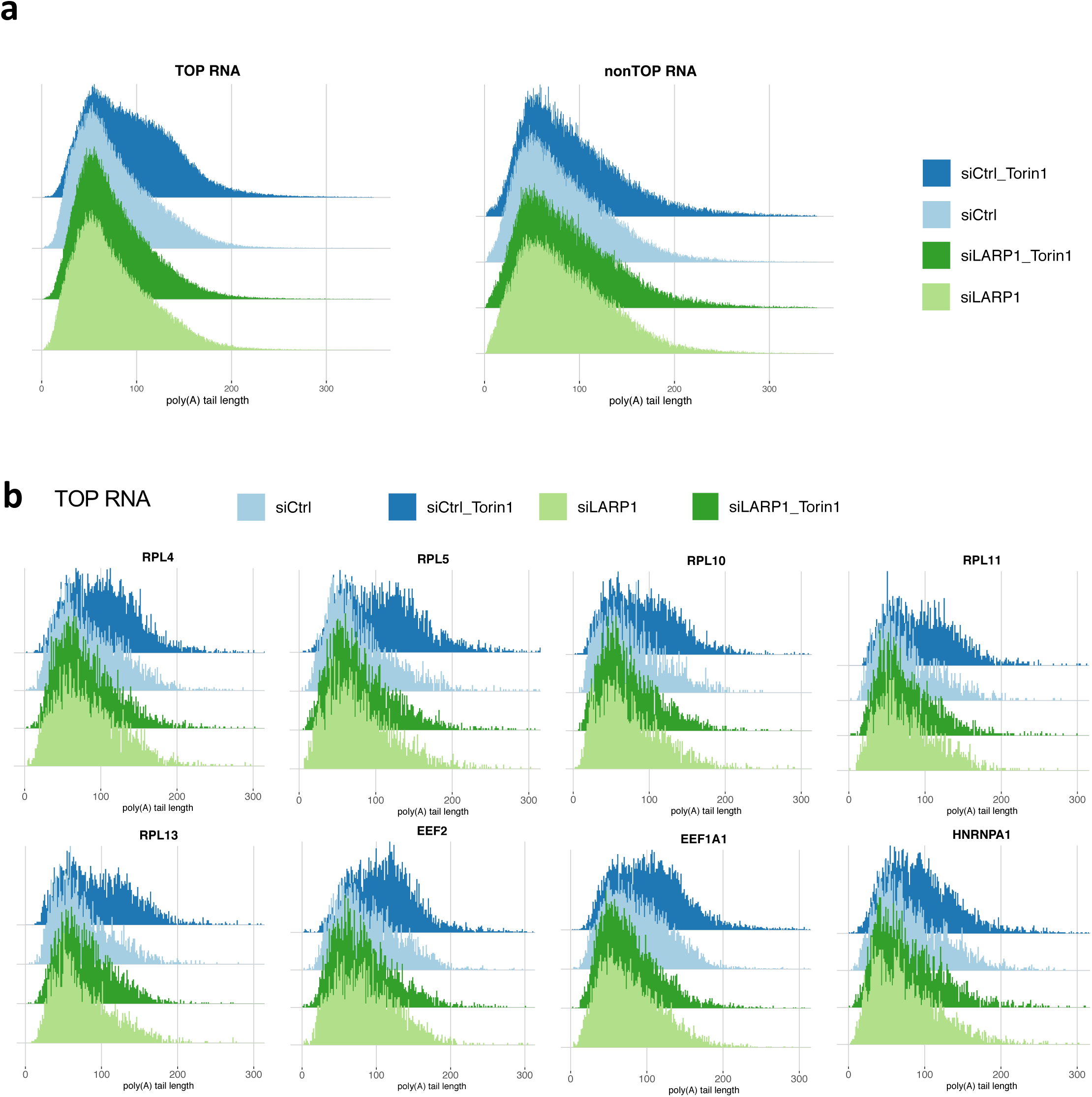
Comprehensive poly(A) tail analysis using nanopore direct RNA sequencing. **a**, Schematic of our approach to prepare the intact mRNA library for dRNA-seq. **b, c**, Density plots showing the distributions of poly(A) tail length of the indicated transcript groups (b) and representative individual TOP transcripts (c).

We next aimed to classify our transcriptome data based on tail length changes. To this end, we define a metric termed PASS (<u>p</u>oly(<u>A</u>) <u>s</u>hift <u>s</u>core) that computes the one-way distance of a cumulative curve of tail length distribution from a siCtrl curve (Fig. 3). The representative results for the individual transcripts are shown (Fig. 3a and Supplementary Fig. 3). PASS analysis of individual genes with > 400 read counts revealed that tail length of TOP mRNAs is mostly lengthened by Torin1 treatment in a LARP1-dependent manner (Fig. 3b), consistently with the results above. To better gain insights into the tail length changes, we applied k-mean (k=8) clustering to the PASS data (Fig. 3c). Large majority of TOP mRNAs were distributed in cluster 1-4 where in many cases both tail elongation after Torin1 treatment and its repression by LARP1 KD were observed. Among the mRNAs exhibiting LARP1-dependent poly(A) lengthening by Torin1, we found that COX7C, RAN, TOMM20 and YWHAZ mRNAs have a sequence that perfectly satisfy the definition of 5’TOP motifs (an invariable cytidine at the cap followed by at least 4 uninterrupted pyrimidines, Supplementary data Fig. 4) (Fig. 3d, depicted as “putative TOP”), and some other mRNAs possess pyrimidine rich sequences at the 5’ ends (Supplementary data Fig. 4). Taken together, our dRNA-seq analysis indicates that TOP mRNA is selectively polyadenylated when mTOR is inhibited and that LARP1 is essential for the process.

**Figure 3.**
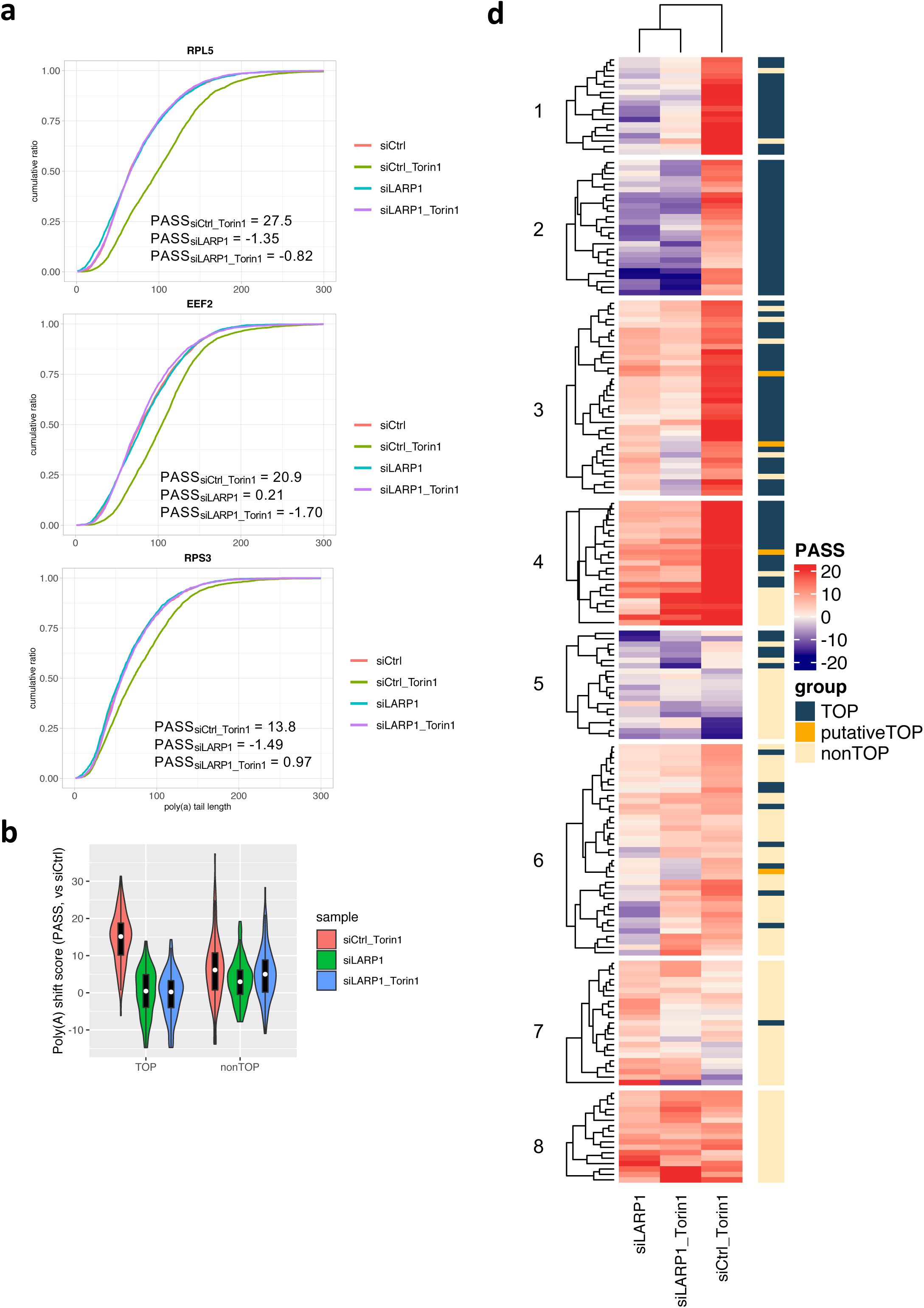
Selective and LARP1-dependent tail lengthening of TOP mRNA. **a**, A metric, termed poly(A) shift score (PASS), comparing the one-way distance of two cumulative curves. **b**, Representative cumulative curves of TOP mRNAs. PASS scores were computed for each mRNA. **c**, Violin and box plots showing the distribution of PASS values for TOP and nonTOP RNAs with > 400 counts (*n* = 102 and 99, respectively). The white dots denote the median; boxes denote the 25th and 75th percentiles; bars denote the range. **d**, Heatmap visualization and *k-means* clustering (k = 8) based on the computed PASS values for the transcripts with > 400 counts (*n* = 201).

**Figure 4.**
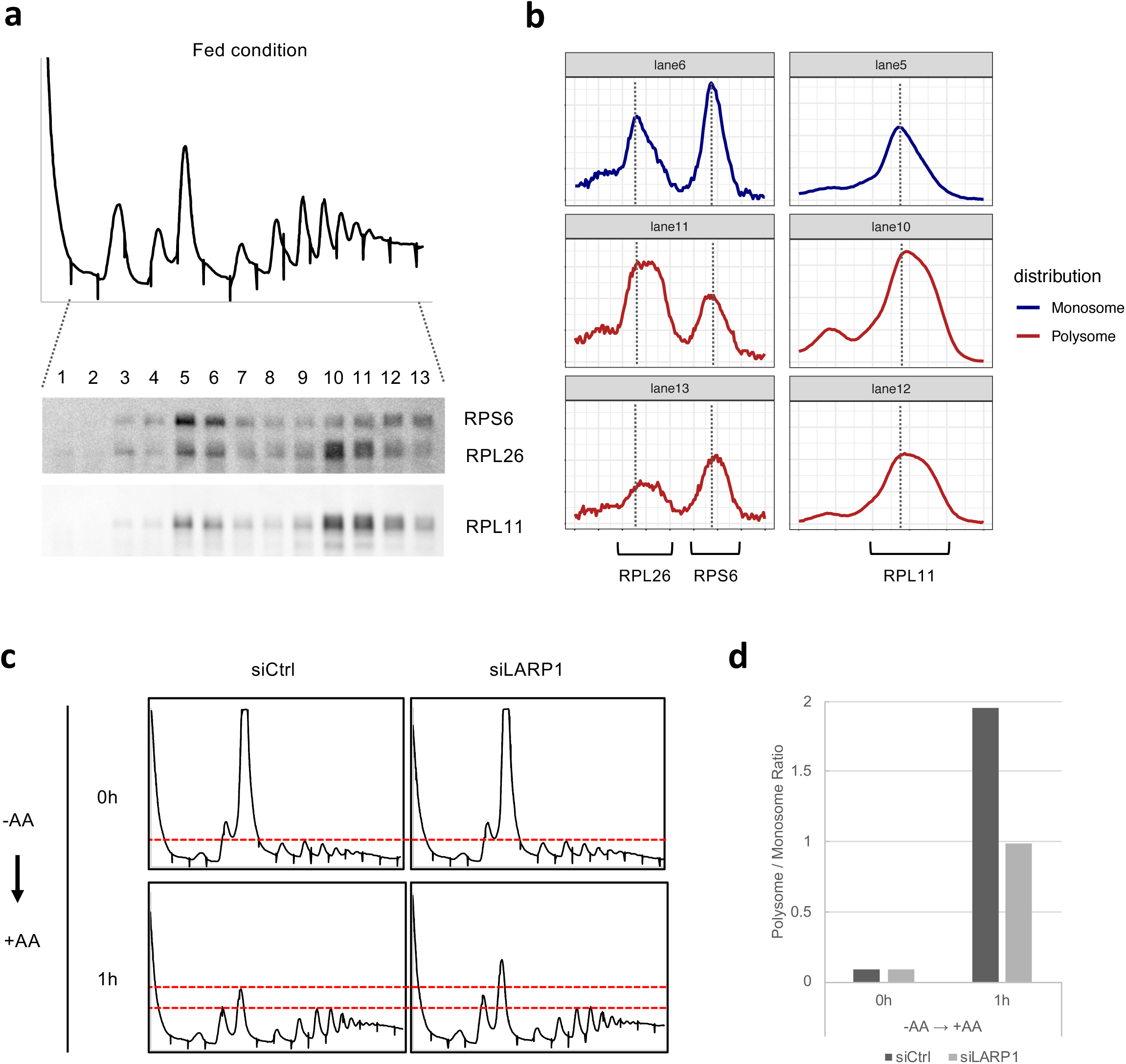
LARP1 ensures quick resumption of translation after amino acid refeeding. **a**, Polysome fractionation using extracts from HEK293T cells cultured under fed condition. The chart shows the absorbance at 254 nm, and northern blotting of the indicated mRNAs in each fraction is shown. **b**, Profiles of the band intensities determined from the indicated lanes in **a. c**, Charts showing the absorbance at 254 nm of polysome fractionation using extracts from siRNA-transfected HEK293T cells cultured under AAS for 14 hrs and re-fed for an hour. **d**, Polysome/monosome ratio was determined by estimating the areas corresponding to monosomal and polysomal fractions.

We compared the changes in the mRNA levels using dRNA-seq data (Supplementary Fig. 5). Considering the differences in the obtained read counts among samples, we computed the KRPG (10^3^ (<u>k</u>ilo) × reads <u>p</u>er <u>g</u>ene) by normalizing read counts for each gene by the total read number (10^3^×Reads of a gene divided by the total read number). Scatter plots of the expression profiles exhibit good correlations among the samples (Supplementary Fig.5a). There are tendencies that Torin1 treatment resulted in slight upregulation of TOP mRNAs, and in contrast, downregulation of nonTOP mRNAs (Supplementary Fig.5b). Besides, TOP mRNA levels were decreased by LARP1 KD, likely indicating RNA destabilization as has been suggested (Supplementary Fig.5b)^3-5^. LARP1 KD did not affect nonTOP RNA abundance. We also estimated the percentage of TOP mRNA read counts in each sample. TOP mRNAs account for 20% of the total cellular mRNA, which is admirably consistent with the earlier estimate (Supplementary Fig. 5c)^15^. The fraction of TOP mRNA in the transcriptome is slightly increased by Torin1 treatment but decreased by LARP1 KD, further suggesting the selective control of TOP mRNA stability (Supplementary Fig. 5c).

We lastly evaluated if the tail elongation boosts recovery of translation from severe repression during AAS when amino acids are replenished. Polysome fractionation markedly revealed that TOP mRNAs with a longer poly(A) are exclusively distributed to the heavy polysomal fractions under fed condition, whereas considerable fraction of the short-tailed counterparts reside in monosomal fractions, suggestive of a positive effect of long poly(A) tail on TOP mRNA translation efficiency (Fig. 4a, b). As TOP mRNAs encode factors essential for translation, we suspected that reduction of poly(A) tail length as well as abundance of TOP mRNA under LARP1 depletion might cause deficiency in translation recovery after refeeding. Indeed, we observed the delayed recovery of the overall translational activity by LARP1 KD (Fig. 4c, d).

Overall, we reveal a dynamic LARP1-mediated poly(A) tail regulation of TOP mRNA responding to amino acid levels. Translation factors are in high demand right after the release from AAS to restore overall protein levels, and thus TOP proteins must be produced with the highest priority. LARP1-dependent preservation of long poly(A)-tailed TOP mRNAs is beneficial for this purpose in both quantitative (increased mRNA stability) and qualitative (increased translational activity by maintaining a long poly(A) tail) aspects. Together, we propose a novel AAS response that ensures full and quick recovery from hostile long-term AAS.

## Acknowledgements

We thank Jared Simpson and Paul Tang (the Ontario Institute for Cancer Research and the University of Toronto) for the supports for the nanopolish polya software. This work was supported by JSPS KAKENHI JP17H03635. We acknowledge the assistance of the Research Equipment Sharing Center at the Nagoya City University. We apologize to all colleagues whose work could not be cited due to space limitations.

## Author Contributions

K.O. contributed to experimental design, computational analysis, interpretation, execution, and manuscript writing and editing. Y. O. performed sample preparation and run of the direct RNA sequencing. K. S. and T. N. performed experiments in Fig.1 and Fig.4. S. H. supervised the project and contributed to experimental design, interpretation, and manuscript writing and editing.

